# Effortful control and cortical brain structure in 5-year-old children: findings from the FinnBrain birth cohort study

**DOI:** 10.1101/2024.12.21.629905

**Authors:** Meri Frantti, Jetro J. Tuulari, Saara Nolvi, Elisabeth Nordenswan, Anni Copeland, Venla Kumpulainen, Eero Silver, Harri Merisaari, Ekaterina Saukko, Eeva-Leena Kataja, Riikka Korja, Linnea Karlsson, Hasse Karlsson, Elmo P. Pulli

## Abstract

The aim of this study was to explore the associations between an aspect of self-regulation (SR), effortful control (EC), and cortical brain structure in 5-year-old children. Efficient EC is a predictor of many attributes and important outcomes in life, such as social-emotional functioning, psychiatric and somatic health, finance, and criminal activity. The early brain correlates of EC are not widely studied, while a better understanding of them would aid in recognizing individuals at risk for maladaptive outcomes. Participants (N = 155) were a part of the FinnBrain Birth Cohort Study in Finland. T1-weighted brain magnetic resonance images were processed using FreeSurfer. The data was statistically analysed with a vertex-wise general linear model. At the age of 5 years, EC and its subscales, Attentional focusing, Inhibitory control (INH), Low intensity pleasure (LIP) and perceptual sensitivity, were assessed via parental report using The Children’s Behaviour Questionnaire. We found positive associations between overall EC and cortical volume in the left superior parietal region and in the right inferior temporal region. We also found positive associations between EC and surface area on the left hemisphere in the superior parietal region. The findings were driven by the EC subscales of INH and LIP with INH linking positively with left surface area and volume, and LIP linking with left cortical volume. We extended the previous literature by shedding light on early structural brain correlates of EC in a large sample of typically developing 5-year-olds. The results differed significantly from previous findings in older children, highlighting the need for longitudinal studies to better understand the neural underpinnings of SR throughout development.

## Introduction

Self-regulation (SR) refers to the capacity to modify one’s behaviour, emotion and cognition in accordance with the environmental framework (Cole et al., 2019; Nigg, 2017). It affects actions and emotions through contributing to the ability to e.g. recognize, express and suppress them in socially adaptive manner. SR can also be considered a part of personality that consists of innate characteristics and is shaped by the experiences in life.

One way to approach SR in young children is to study temperament traits (Goldsmith et al., 1987). Rothbart’s psychobiological theory of temperament conceptualizes temperament in young children as the individual differences in reactivity and regulation, with reactivity divided into positive and negative emotional reactivity, and aspects of regulation usually referred to as effortful control (EC). In this study, we focus on EC, which represents the ability to control one’s actions under conflict, plan the future actions and to detect errors (Rothbart, 2007; Nigg, 2017). Thus, EC can be considered a trait-level aspect of top-down SR (Nigg, 2017). The precursors of EC are observed already in infancy (Eisenberg et al., 2014), but moderate stability of EC is reached from toddlerhood or preschool onwards (Rothbart, 2007). Childhood EC also predicts adulthood conscientiousness, a personality trait linked with SR and favourable life outcomes (Eisenberg et al., 2014; Pérez-Edgar & Fox, 2005; Rothbart, 2007).

EC and other aspects of SR are important predictors for a myriad of later life outcomes: higher self-control in childhood is positively linked to social skills and academic achievement and negatively linked to aggressive and criminal behaviour, mental and physical health issues, obesity and cigarette smoking (Robson et al., 2020). Lower EC during adolescence is also linked to the risk of subsequent substance use disorder (Cheetham et al., 2017; Robson et al., 2020). In addition, higher EC at 2–8 years was associated with more prosocial behaviour and less internalizing and externalizing symptoms (Robson et al., 2020; Slobodskaya et al., 2020). A better understanding of the early underlying mechanisms of EC could help identify individuals at risk and create interventions to prevent unfavorable outcomes later in life.

Particularly, the knowledge of early neurobiological factors underlying EC could be of interest in understanding the dynamic development of SR and EC. Traditionally, as reviewed by Kelley et al., 2015 and Fiske & Holmboe, (2019), the main neural areas linked to SR are thought to be the prefrontal cortex (PFC), particularly the ventromedial and the lateral regions, and the anterior cingulate cortex (ACC). Currently there are few child or adolescent magnetic resonance imaging (MRI) studies that used questionnaire data investigating SR in real-life contexts (see Table 1). Out of the 20 studies found, only 2 included both MRI and SR assessment questionnaire of children under 6 years of age. Hadaya et al. (2023) conducted a study where children born preterm had an MRI scan at 38–53 weeks post-menstrual age and a neuropsychological follow up at 4–7 years. Interestingly, the subgroup that had highest EC scores also displayed larger relative volumes in the left insula and bilateral OFC and higher degree centrality in an overlapping region in the lOFC. Vanes et al. (2021) also focused on preterm children and studied the associations between neonatal brain structure, home environment and childhood outcomes with MRI conducted at term and questionnaire data gathered at the age of 4–7 years. They found smaller fronto-insular volume in a cluster of participants characterized by executive deficits, attention deficit hyperactivity disorder symptoms and autism spectrum symptoms (the result did not survive correction for multiple comparisons). Notably, the neuroanatomical findings in studies with younger participants differ from those usually seen in older participants (Kelley et al., 2015) and research on special population such as preterm born children should be complemented by research in general population participants.

**Table 1:**
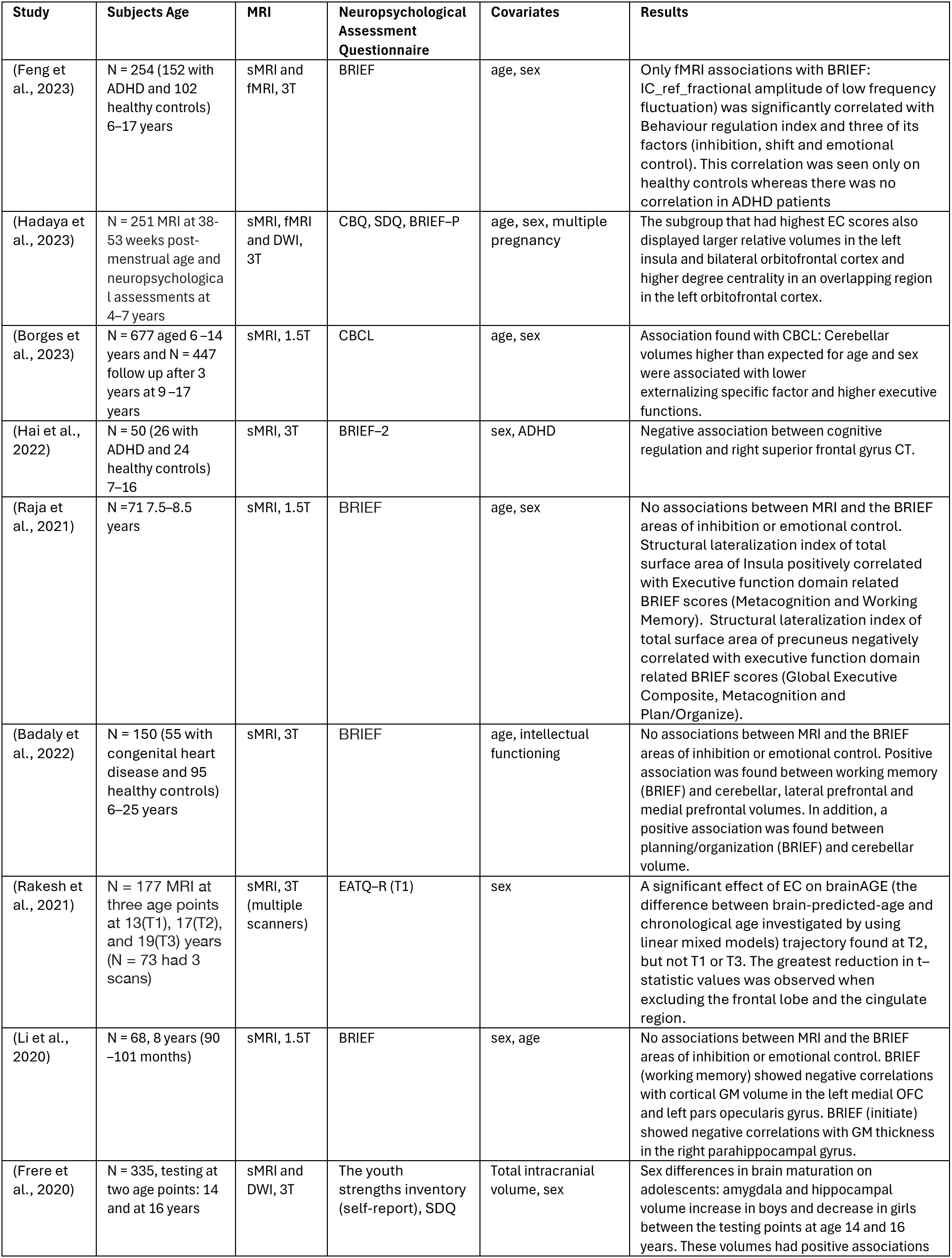
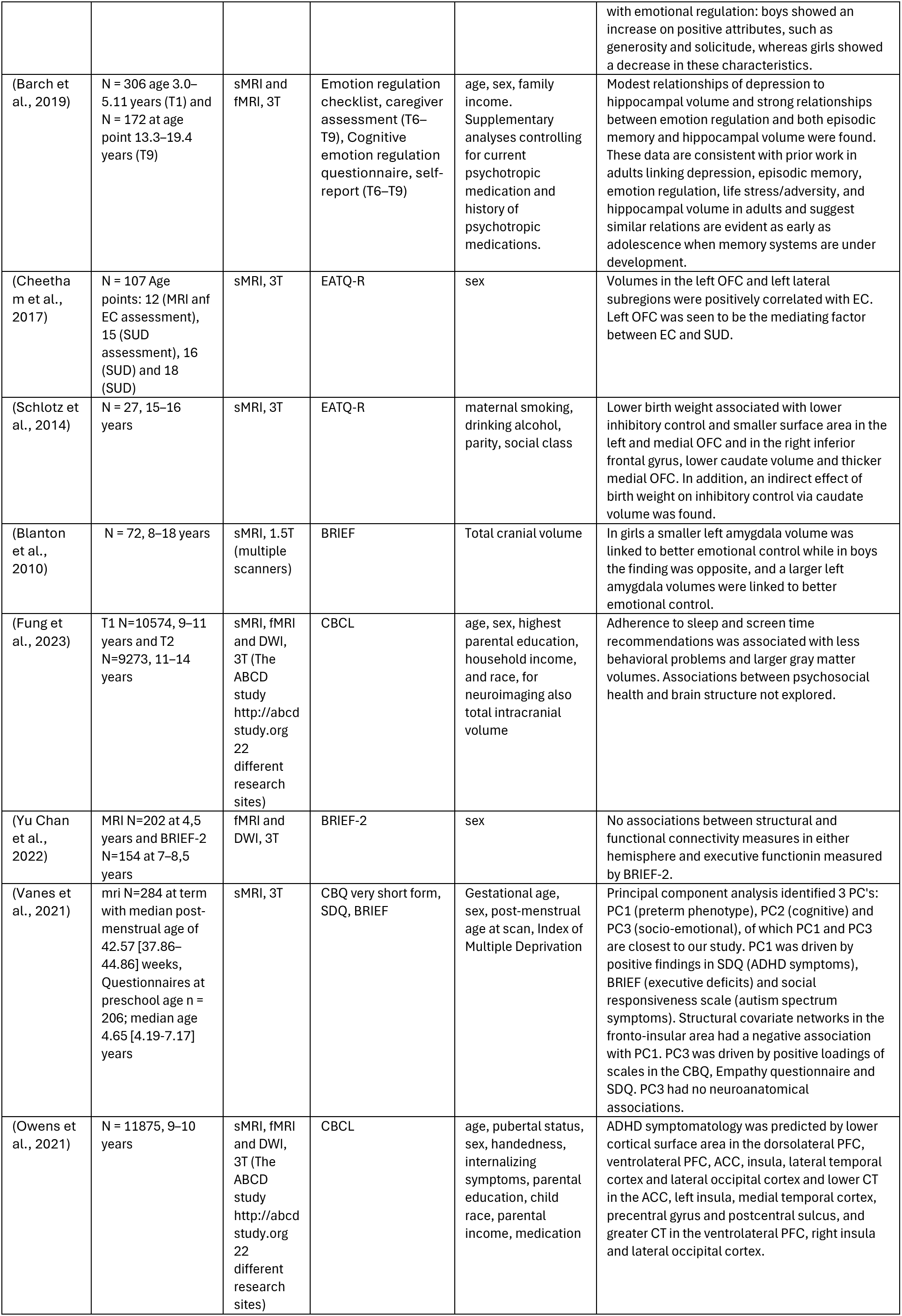
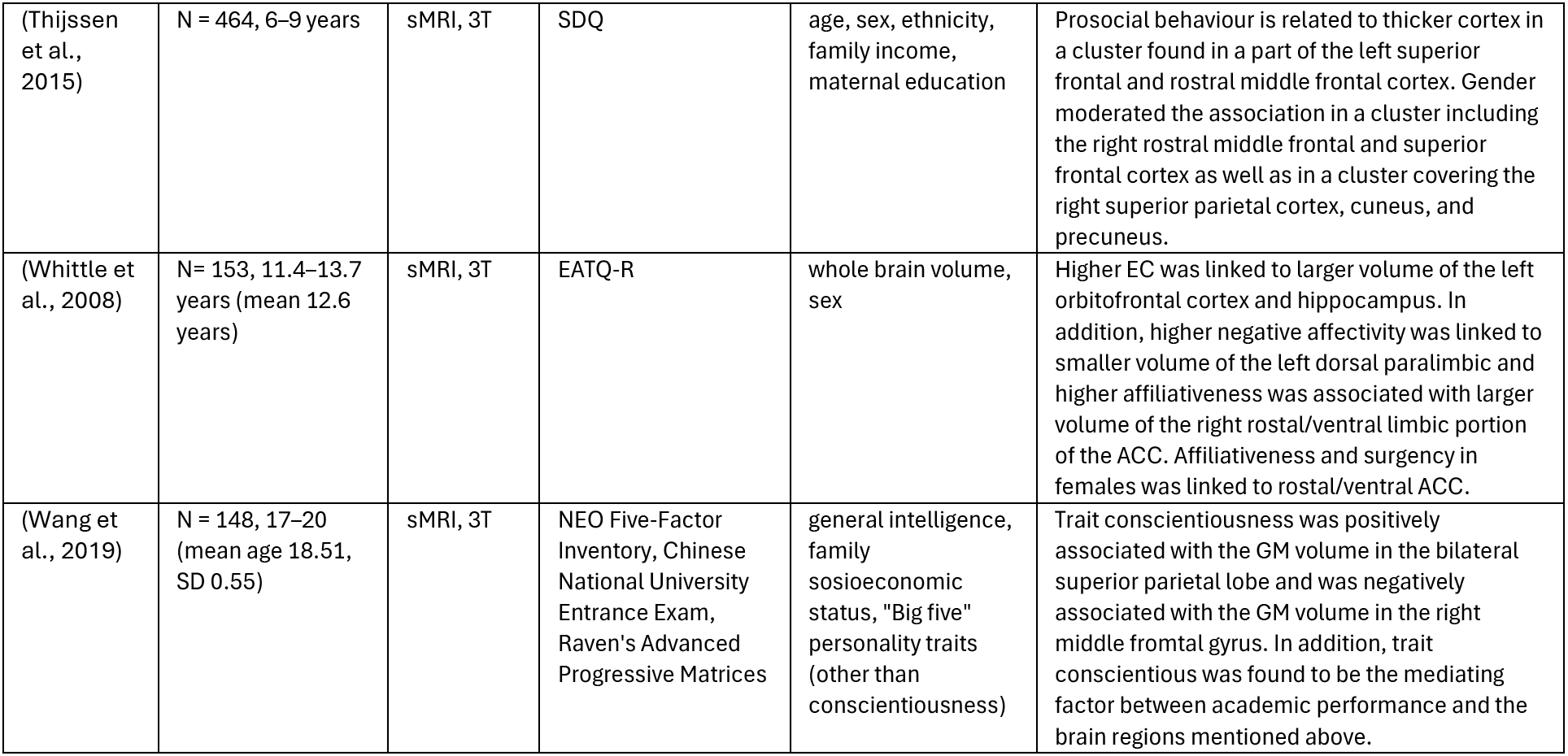
We conducted a Pubmed search on January 21^st^ 2024 with the following query: mri AND (structur* OR volume OR area OR "cortical thickness") AND (infan* OR toddler OR child OR childre* OR adolesc*) AND (self-control OR "effortful control" OR "executive function" OR "inhibitory control") and the last two articles on the table are added from our previous searches. BRIEF = Behaviour rating inventory of executive function (parental asessment) ADHD = Attention Deficit Hyperactivity Disorder CBQ = Child behavioral questionnaire SDQ = Strengths and difficulties questionnaire (self-report) CBCL = Child behaviour checklist EATQ-R = Early Adolescent Temperament Questionnaire-Revised SUD = Substance use disorder PFC = prefrontal cortex ACC = anterior cingulate cortex

To summarize, the research on early childhood neural underpinnings of EC, an aspect of SR, remains scarce. To the best of our knowledge, this is the first large cross-sectional study examining EC and brain structure in preschool age general population children. Some earlier studies have examined this topic in a sample with a wider age range (Badaly et al., 2022; Feng et al., 2023). As both the brain and EC are still in a developing state at 5 years of age, the neural basis of EC could be substantially different from that in older participants (Hadaya et al., 2023). Furthermore, grey matter (GM) volume reaches its peak at 6 years of age (Bethlehem et al., 2022; Courchesne et al., 2000) and then starts to decrease (Bethlehem et al., 2022). There is regional variation in the age of peak GM volumes, ranging from age 2 to age 10 years (Bethlehem et al., 2022). Therefore, if better EC is associated with either faster or slower development in certain brain regions, the direction of those associations changes in preschool or early school age, complicating the interpretation of results from studies with wide age ranges.

The aim of this study was to address the gaps in the previous literature by utilizing a large cross-sectional sample of typically developing 5-year-old children to examine the neural bases of EC, at an age when both brain and EC are still dynamically developing. This study will provide novel information on the structural associations of EC at this developmentally crucial age. We did not set an explicit hypothesis for this study, because there is no previous structural MRI research on EC in this specific age group.

## Methods

This study complies with the Declaration of Helsinki and was approved by the Joint Ethics Committee of the University of Turku and the Hospital District of Southwest Finland. ETMK: 26/1801/2015 for the neuropsychological measurements, and ETMK 31/180/2011 for the neuroimaging.

### Participants

Participants were a part of the population-based FinnBrain Birth Cohort Study (www.finnbrain.fi), which is a prospective pregnancy cohort aiming to explore prenatal and early life stress and their effect on child brain development and health (Karlsson et al., 2018). The families were recruited during the mothers’ first trimester ultrasound at gestational week (GW) 12, between December 2011 and April 2015 in the South-Western Hospital District and Åland Islands in Finland. In addition to an ultrasound-verified pregnancy, sufficient knowledge of the Finnish or Swedish language was required for participation. A written informed consent was provided by the parents prior to the children’s study visits.

The neurocognitive assessments were conducted at the age of 5 years between October 2017 and December 2020. Total of 1288 families were contacted, from which 974 (75.6%) were reached by telephone and 545 (42.3%) ended up participating in a study visit (304 boys (55.8%), mean age 5.0 (SD 0.1), range 4.9–5.4 years). Mothers of the children who did not participate in the neurocognitive visit (out of the 1288 contacted families) had lower education level (χ^2^ (2) = 30.94, p < 0.001), a lower monthly income (χ^2^ (3) = 11.65, p = 0.009) and were younger (t(1286) = -4.130, p < 0.001) compared to the mothers in the families that participated in the neurocognitive visits.

Next, the families that had attended the neurocognitive visit were prioritized in recruitment for the neuroimaging visits, where 541 families were contacted and 478 (88.4%) of them were reached. In total, 203 (37.5%) participants attended imaging visits (111 boys, 92 girls, mean age 5.4 (SD 0.1), range 5.1–5.8 years). Participants were healthy, typically developing children and 182 were right-handed, 14 left-handed and 7 had no preference. 196 participants attended both, neurocognitive and neuroimaging study visits. Mothers of the children who participated in the neuropsychological visits but not in the neuroimaging visits were older (t (369) = 1.97, p = .047) but did not differ in education level or monthly income compared to the mothers in the families that participated in the MRI visit.

The original goal was to conduct all the neuroimaging study visits when the subjects were between the ages 5 years 3 months and 5 years 5 months, however some of the participants ended up being older than planned due to a break in our study visits caused by the COVID-19 pandemic (152/203 (74.9%) of the participants attended the visit within the intended age range).

The exclusion criteria for the neuroimaging study were: 1) born before GW 35 (before GW 32 for those with exposure to maternal prenatal synthetic glucocorticoid treatment) 2) developmental anomaly or abnormalities in senses or communication (e.g., blindness, deafness, congenital heart disease) 3) known long-term medical diagnosis (e.g., epilepsy, autism) 4) ongoing medical examinations or clinical follow up in a hospital (meaning there has been a referral from primary care setting to special health care) 5) child use of continuous, daily medication (including per oral medications, topical creams and inhalants, apart from desmopressin (®Minirin) medication, which was allowed) 6) history of head trauma (defined as concussion necessitating clinical follow up in a health care setting or worse) 7) metallic (golden) ear tubes (to assure good-quality scans) and routine MRI contraindications Out of the total 196 MR images, 173 were of adequate quality. Of these subjects, 18 parent reports of EC were missing, resulting in a final sample size of 155. The demographic information of the participants is presented in Table 2.

**Table 2.**
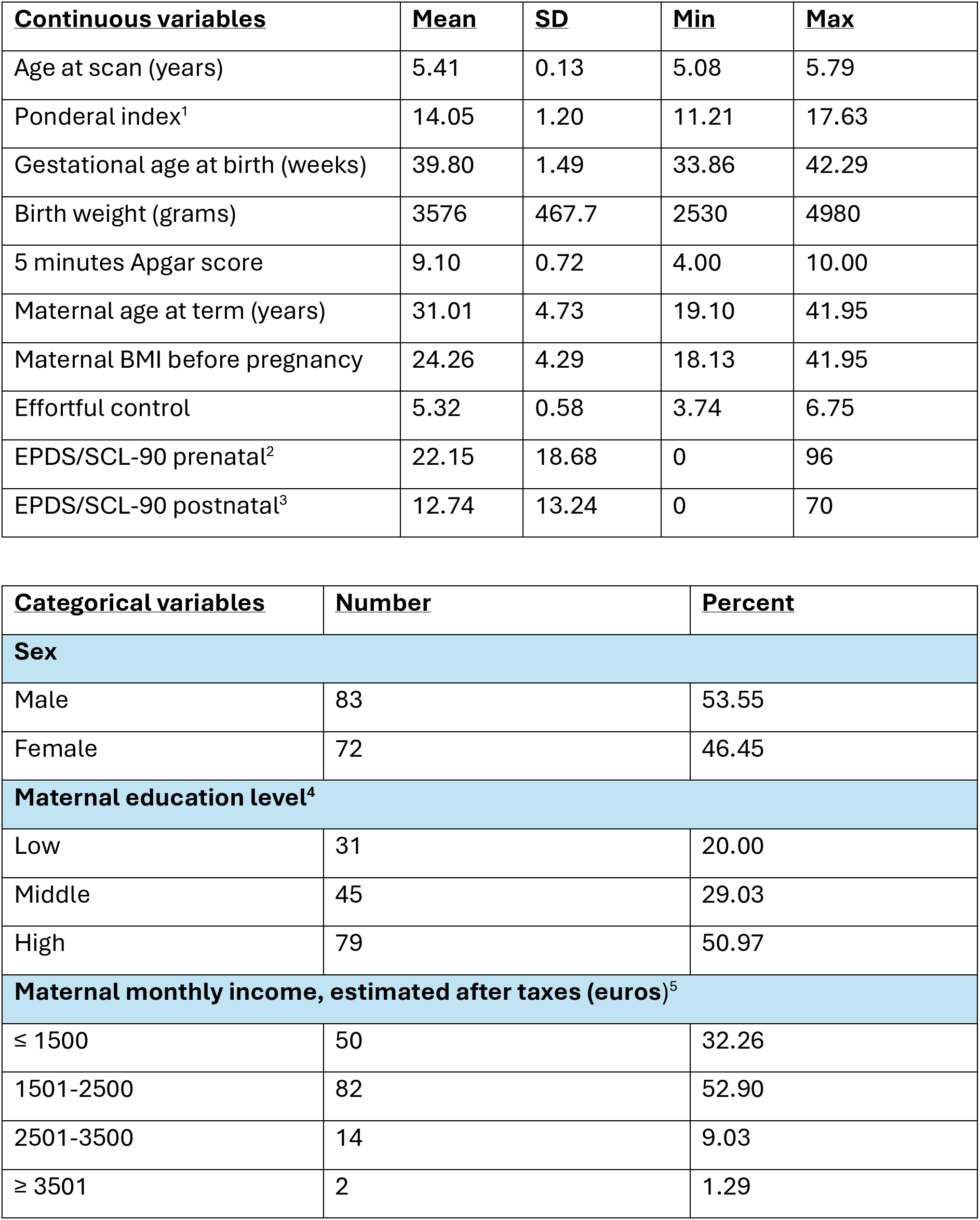
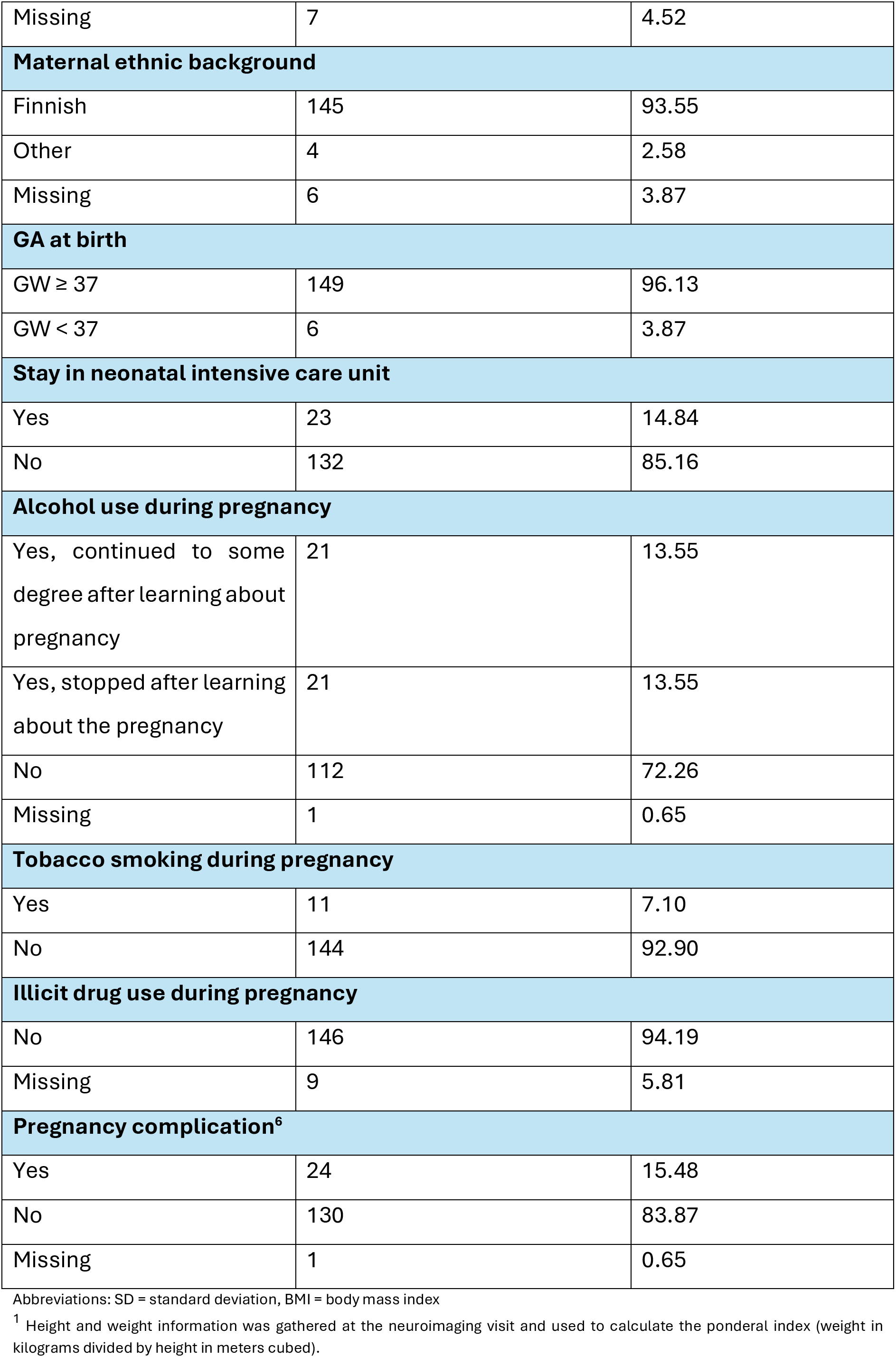
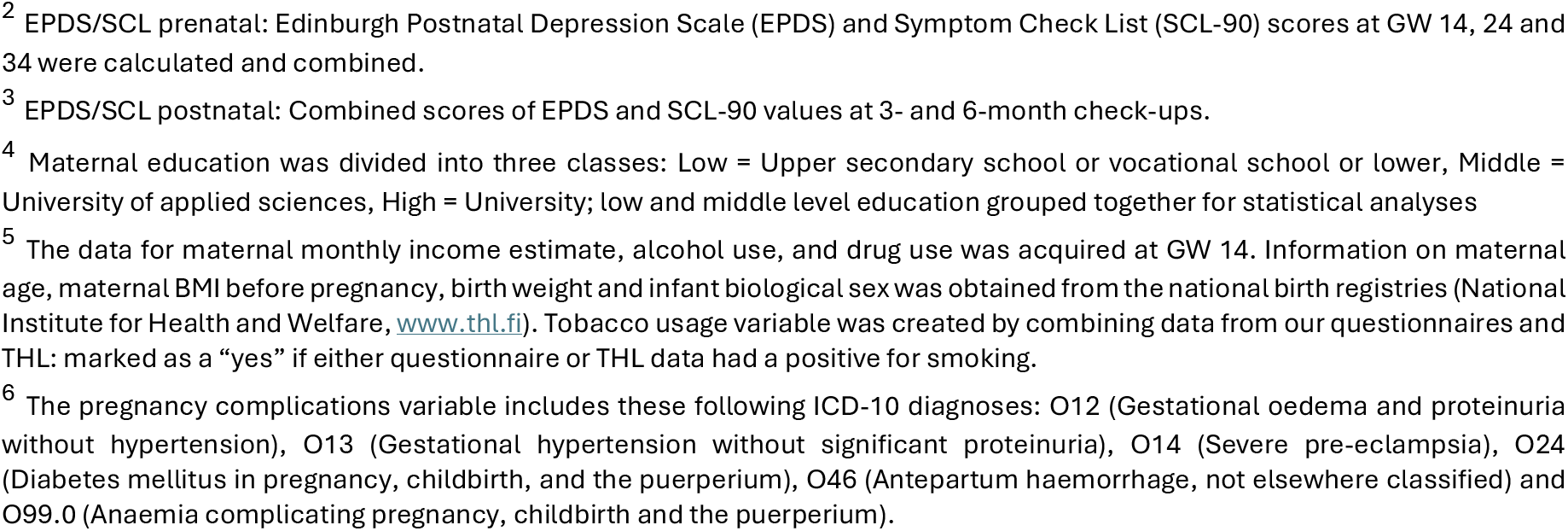
Participant demographics and maternal medical history variables (N= 155)

### Procedures

The parents reported about child temperament by responding to the Children’s Behaviour Questionnaire (CBQ) at 5 years of age during the FinnBrain Child Development and Parenting Functioning Lab’s neurocognitive study visit. After that visit, the participating families were invited to the MRI visit, where structural T1-weighted images were collected as a part of a max. 60-minute scan. The study report follows the Strenghtening the Reporting of Observational Studies in Epidemiology (STROBE) guidelines (Vandenbroucke et al., 2007), see Supplementary File 1 for the attached checklist.

### Neurocognitive study visits

The neurocognitive study visits for 5-year-old children included neurocognitive testing, eye-movement tracking, mother-child interaction assessment and questionnaires filled out by the parents, including CBQ. Child EC was assessed with parental reports at the age of 5 via CBQ short form, a revised 94-item version of the more extensive 195-item measure (Rothbart et al., 2001). CBQ is a caregiver report measure designed to provide a detailed assessment of temperament in children 3 to 7 years of age (Rothbart et al., 2001). The parents responded the CBQ by evaluating their child’s behaviour during the past six months on a Likert scale from 1–7 (1 being “not at all” and 7 being “yes, most of the time”), higher score indicating higher level of certain characteristics. The CBQ includes 15 dimensions of temperament forming the following main dimensions: EC, negative affectivity and surgency. Our main response variable in this study, EC, consists of 4 subscales: 1) Attentional focusing (ATTFO), the tendency to maintain attentional focus when performing a task; 2) Inhibitory control (INH), the ability to plan and suppress certain approach responses under instructions; 3) Low intensity pleasure (LIP), the pleasure experienced in a situation involving low-intensity stimuli; and 4) Perceptual sensitivity (PER), the detection of low-intensity stimuli. The statements considering the 4 subscales of EC were combined, coming to a total of 25 items considering EC and the average was calculated.

EC main dimensions showed good internal consistency, with Cronbach’s alpha 0.829. Cronbach’s alpha for the subscales ranged from 0.644 to 0.787, which was considered adequate for the exploratory analyses. One missing answer was allowed (per factor), and the missing one was imputed with the mean of other answers in the EC questions.

We did not gather data on who filled out the CBQ form, however the Child Behaviour Checklist (CBCL) form was filled during the same neurocognitive study visit and included information about the filling parent. Assuming almost complete overlap between CBQ and CBCL fillers, most of the CBCL fillers were mothers (92.90%). In some cases, the filling parent was the dad (3.9%), or the parents filled out the form together (1.30%), with data missing from 1.90% of the cases.

### Neuroimaging study visits

The study visit protocol is described in more depth in previous work from FinnBrain (Copeland et al., 2021; Pulli et al., 2022). Practice materials were delivered to each family prior the study visit. At the beginning of the study visit a written informed consent from both parents and a verbal agreement from the child was attained. In the course of the two-hour preparation time before the scan the child got familiar with the staff, went through a practice scan, and enjoyed a light meal. During the scan the participants were awake or in natural sleep. Apart from the fMRI sequence, the participants could watch a movie or a cartoon of their choice if they wanted to. A parent and a research staff member were always present in the scanning room throughout the scan. Everyone in the room had their hearing protected with both earplugs and headphones.

### MRI data acquisition

Siemens Magnetom Skyra fit 3T with a 20-element head/neck matrix coil was used in scans. Generalized Autocalibrating Partially Parallel Acquisition (GRAPPA) technique was used to accelerate image acquisition (parallel acquisition technique [PAT] factor of 2 was used). The max. 60-minute scan protocol included a high resolution T1 magnetization prepared rapid gradient echo (MPRAGE), a T2 turbo spin echo (TSE), a 7-minute resting state functional MRI and a 96-direction single shell (b = 1000 s/mm^2^) diffusion tensor imaging (DTI) sequence (Kumpulainen et al., 2022; Merisaari et al., 2019; Rosberg et al., 2022) as well as a 31-direction with b = 650 s/mm^2^ and a 80-direction with b = 2000 s/mm^2^. For the purposes of the current study, we acquired high resolution T1-weighted images with the following sequence parameters: TR = 1900 ms, TE = 3.26 ms, TI = 900 ms, flip angle = 9 degrees, voxel size = 1.0 x 1.0 x 1.0 mm^3^, FOV 256 x 256 mm^2^. The scans were planned as per recommendations of the FreeSurfer developers (https://surfer.nmr.mgh.harvard.edu/fswiki/FreeSurferWiki?action=AttachFile&do=get&target=FreeSurfer_Suggested_Morphometry_Protocols.pdf, at the time of writing).

### Image Processing

Cortical reconstruction and volumetric segmentation were performed with the Freesurfer 6.0.0 image analysis suite, available for download online (http://surfer.nmr.mgh.harvard.edu/). The technical details of these procedures are described in prior publications (Dale et al., 1999; Dale & Sereno, 1993; Fischl et al., 2000, 2001, 2002; Fischl, Salat, et al., 2004; Fischl, Sereno, & Dale, 1999; Fischl, Sereno, Tootell, et al., 1999; Fischl, van der Kouwe, et al., 2004; Han et al., 2006; Jovicich et al., 2006; Reuter et al., 2010, 2012; Ségonne et al., 2004).

After the initial FreeSurfer processing, all images were visually checked and manually edited (skull fragments removed if they were influencing the pial border, errors corrected in the border between GM and WM and arteries removed). After the edits, the FreeSurfer recon-all was run once more. For a more detailed description of the image processing procedure, please see our previous article (Pulli et al., 2022).

### Covariate selection for vertex wise statistics

The potential confounders were chosen based on our previous work on 5-year-olds (Silver et al., 2022). Covariates in our analyses were the child’s gender, age at scan, ponderal index (mass in kilograms divided by height in meters cubed; measured during the neuroimaging visit), maternal age at term and maternal education level. Maternal education data was gathered from questionnaire data from GW 14 or 5 years of child age by using the highest degree reported (two classes, university vs. other degree).

We performed sensitivity analyses controlling for the following factors: maternal body mass index (BMI) before pregnancy (Li et al., 2016; Ou et al., 2015) and gestational weeks at birth. In other sensitivity analyses, a part of the sample was excluded: we took into account alcohol exposure in utero (missing data interpreted as no exposure) (Donald et al., 2015), tobacco exposure (Chang et al., 2016; Knickmeyer et al., 2016) and children with neonatal intensive care unit (NICU) stay were excluded (Aoki et al., 2020). We also controlled for maternal scores from the Edinburgh Postnatal Depression Scale (EPDS), a self-report questionnaire of depressive symptoms within the past 7 days (Cox et al., 1987), and Symptom Checklist 90 (SCL-90), an assessment of psychiatric symptoms within the past month (Derogatis, 1994). We calculated prenatal score combining EPDS and SCL-90 scores from GW 14, 24 and 34. Postnatal value consisted of EPDS and SCL-90 values at 3- and 6-monts of child age. The EPDS consists of 10 items considering 3 subscales: anxiety, anhedonia and depression, and the total score ranges from 0 to 30, larger score meaning more depressive symptoms. We used the Anxiety Subscale of the SCL-90, score ranging from 0–40, higher score meaning more severe symptoms.

### Statistics

IBM SPSS for Mac, version 27.0 (IBM Corp., Armonk, NY, USA) was used for the statistical analyses concerning demographics.

As instructed in the FreeSurfer manual, we pre-smoothed fsaverage surfaces for analyses with Query, Design, Estimate, Contrast (Qdec), a single-binary application included in the FreeSurfer software suite (http://surfer.nmr.mgh.harvard.edu/). Qdec is a graphical user interface for a statistics engine running a vertex-by-vertex general linear model (GLM). For display purposes, we used the standard FreeSurfer’s fsaverage in MNI305 space (MNI = Montreal Neurological Institute).

We tested for clusters with statistically significant associations between EC and cortical GM volume, SA and CT. In addition to EC, we performed post hoc analysis on all its subscales (ATTFO, INH, LIP and PER).

The data was smoothed with a kernel of 10 mm full width at half maximum. A Monte Carlo Null-Z Simulation was run with a z-value threshold of 1.3, corresponding to p = 0.05 (Hagler et al., 2006). For performed sensitivity analyses, please see “Covariate selection for vertex wise statistics”. Age at scan was squared for the purposes of running Qdec.

## Results

### EC and brain volume

Figure 1 presents the associations between brain volume and EC. The significant associations were positive. On the left hemisphere there was a cluster in the supramarginal region (peak p < 0.0001, size: 1961.59 mm^2^, peak coordinates: (-49.6, - 48.0, 44.6)). On the right hemisphere there was a cluster in the inferior temporal region (peak p = 0.0025, size: 1253.93 mm^2^, peak coordinates: (46.9, -21.1, -26.6)).

**Figure 1:**
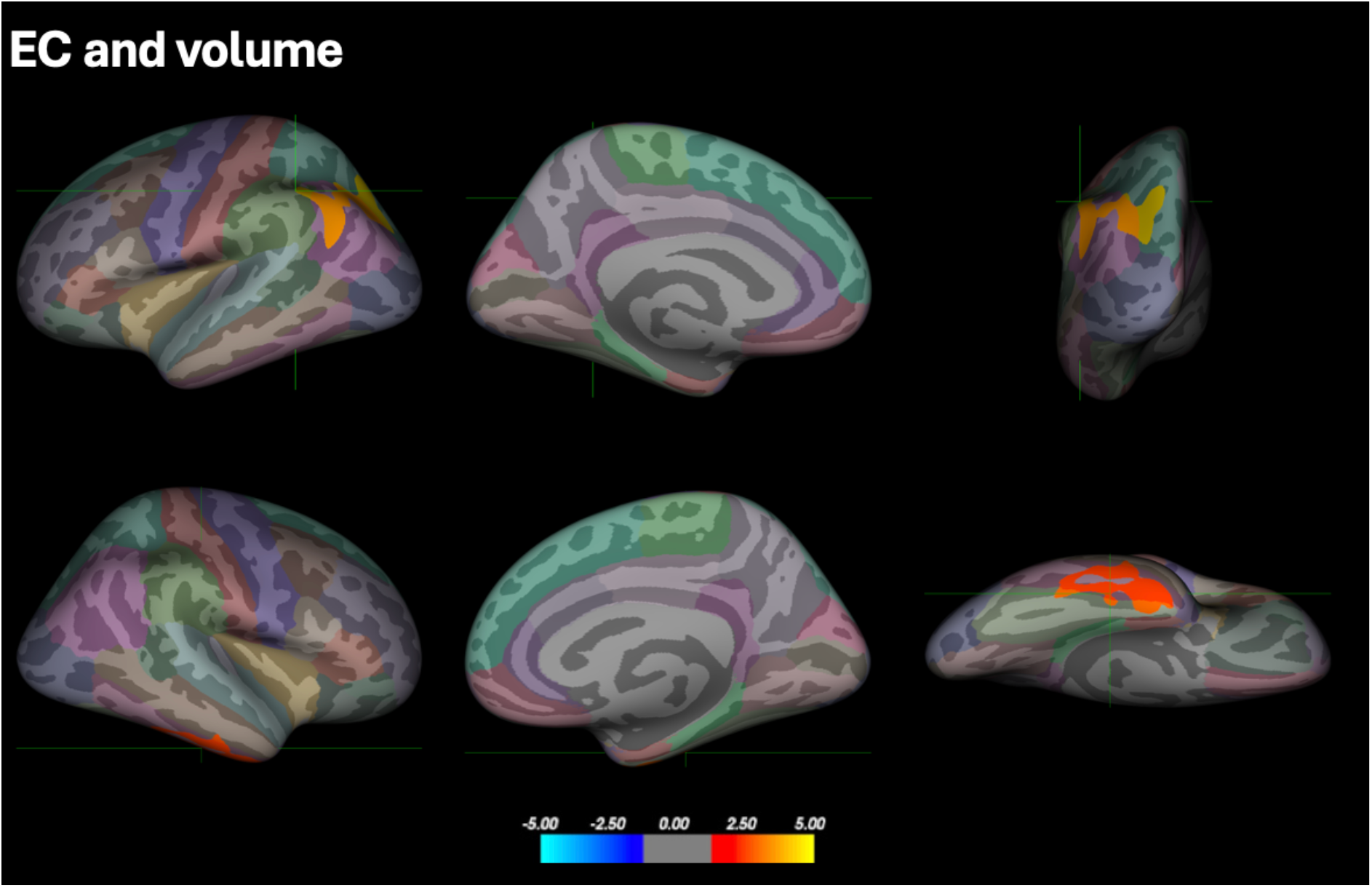
Positive associations between EC and cortical volume. Result corrected for multiple comparisons using Monte-Carlo Null-Z simulation. The position of the green crosshair indicates the most statistically significant vertex in statistically significant clusters. Left hemisphere is pictured on the top row and right hemisphere on the bottom row. From left to right, first is lateral and second medial view. The last pictures on the right, where both clusters are best visible, are posterior view of the left hemisphere and basal view of the right hemisphere. Color coding of regions according to the Desikan-Killiany atlas. Scale: -log_10_(p).

### EC and surface area

The cluster coordinates of the positive association in the left hemisphere were the same between EC and SA, INH and SA, and LIP and volume in the superior parietal region. Figure 2 presents the associations between SA and EC, where the only significant association was positive. The cluster was located on the left hemisphere in the superior parietal region (peak p = 0.0007, size: 1202.95 mm^2^, peak coordinates: (-23.9, -78.5, 16.8)). No clusters were found on the right hemisphere.

**Figure 2:**
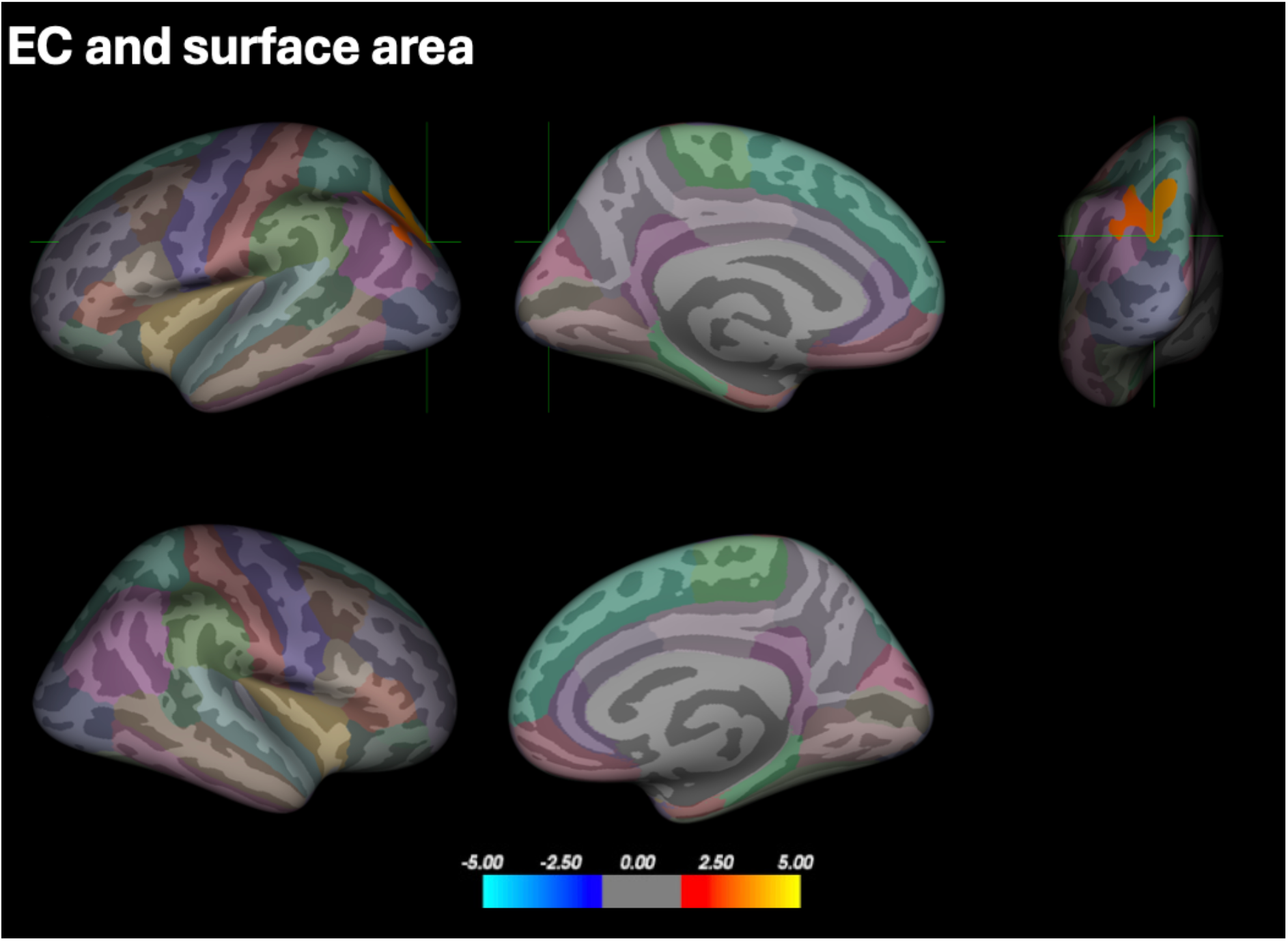
Positive association between EC and SA. Result corrected for multiple comparisons using Monte-Carlo Null-Z simulation. The position of the green crosshair indicates the most statistically significant vertex in statistically significant clusters. Left hemisphere is pictured on the top row and right hemisphere on the bottom row, the first picture from left to right is lateral view and the second is medial view. On the top right corner also a posterior view of the left hemisphere is included. Color coding of regions according to the Desikan-Killiany atlas. Scale: -log_10_(p).

### EC and CT

We also analysed CT on Qdec, using the same variables, but no results were found after conducting the Monte Carlo simulation.

### Post hoc analyses

In the subscales of EC, associations were only found between neural correlates and INH and LIP, all significant associations were positive and only visible on the left hemisphere (Figure 3).

**Figure 3:**
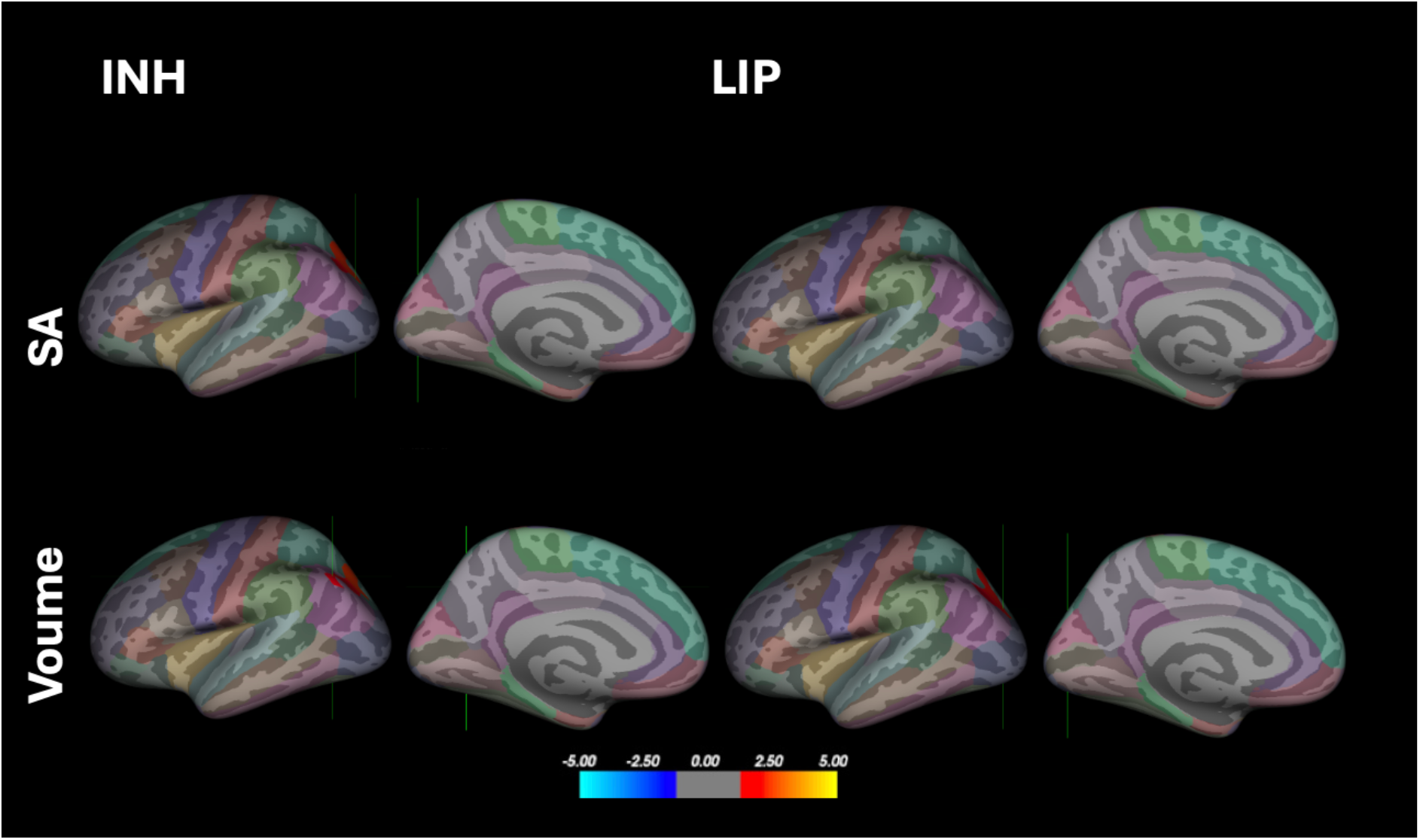
The association between Inhibitory Contro (INH)l and Low Intensity Pleasure(LIP) with SA on the top row and with volume on the bottom row. Both lateral and medial views of the left hemisphere included. The right hemisphere is not included, since no significant associations were found. Result corrected for multiple comparisons using Monte-Carlo Null-Z simulation. The position of the green crosshair indicates the most statistically significant vertex in statistically significant clusters. Color coding of regions according to the Desikan-Killiany atlas. Scale: -log_10_(p).

Cluster representing the association between INH and SA was in the superior parietal region (peak p = 0.0200 size: 786.69 mm^2^, peak coordinates: (-23.9, -78.5, 16.8)). Whereas the cluster representing the association between INH and brain volume was in the inferior parietal region (peak p = 0.0151, size: 989.46 mm^2^, peak coordinates: -37.8, - 67.0, 45.6).

No cluster was found when analysing the association between LIP and SA. The cluster representing the association between LIP and volume was in the superior parietal region (peak p = 0.0417, size: 851.96 mm^2^, peak coordinates: (-23.9, -78.5, 16.8).

### Sensitivity analyses

No major differences were found in the sensitivity analyses. The results are presented in detail in “Supplementary Materials”.

## Discussion

In this study, we investigated the associations between EC and structural cortical brain metrics, more precisely volume, SA and CT, in a large sample of typically developing 5-year-olds. We also examined the subscales of EC and their relation to cortical brain anatomy, to assess whether the effects were driven by any specific subscales. All significant findings were positive and all significant results were found on the left side, except for one. Higher EC (and its subscales) was generally related to higher volume or larger SA in the child brain, but no associations found between EC or any of its subscales and CT.

To summarize the results, higher child EC was associated with a cluster located in the superior parietal, inferior parietal and supramarginal region. At 5-years of age the GM volume is close to reaching its peak, which happens at 5.9 years (Bethlehem et al., 2022; Courchesne et al., 2000). However, the regional peaks of GM volume in the cortical regions vary notably between ages 2–10 years (Bethlehem et al., 2022) and occipito– parietal volumes peak first before the age of 5 years (Aubert-Broche et al., 2013; Bethlehem et al., 2022). The cluster includes regions which peak around or before 5 years of age, making it difficult to conclude whether the children with higher EC are further along the brain development or developing slower. We found a positive correlation between EC and right inferior temporal SA, a region still growing towards its peak (Bethlehem et al., 2022), which suggest that EC is higher in children that are further along the brain development. However, based on a single cross-sectional study we can only speculate about developmental trajectories. Longitudinal studies with repeated imaging that allow comparing the same individuals at different age points close to each other, will be crucial in identifying the neural underpinnings of EC at different stages of development.

Based on post hoc analysis, these associations were driven by EC indicators of Low Intensity Pleasure (LIP) and Inhibitory Control (INH) specifically. Though links between EC and the parietal lobe has not been reported previously, there are reports of higher parietal lobe volume linking with conscientiousness (Wang et al., 2019) which is considered continuum for higher childhood SR (and thus, EC). The parietal lobe is a part of the parieto–frontal integration theory (P–FIT), which has been linked to cognitive ability, see review by Jung & Haier, (2007). Cognitive abilities are connected to executive function, a concept closely related to SR and inhibitory control, see review by Fiske & Holmboe, (2019).

We found no supporting evidence to the links between EC and either PFC or ACC, which are the most important neural correlates indicated in the previous studies (Kelley et al., 2015) and parts of the P-FIT (Jung & Haier, 2007). In our study, none of the results in the OFC survived the Monte Carlo simulation and no negative results survived altogether. Previous MRI studies have found a positive correlation between lOFC volume and EC on 12-year-olds (Whittle et al., 2008). This same correlation was also found in a lesion study on adults by Bechara et al. (2000) and a similar result was found by Hadaya et al. (2023), where on 4–7 -year-olds higher bilateral OFC and left insula volume and centrality in the lOFC was linked to higher EC. Furthermore, like in previous studies, most findings in our study were found in the left hemisphere, but the specific regions differed significantly.

One possible reason for the discrepancies between the previous research and our study may lie in the age differences between the studies. EC has been positively associated to the difference between brain-predicted age (based on cortical and subcortical measures from MRIs) and chronological age in regions of the frontal lobe at the age-point of 16 years, but the association was not significant at other age points, which were at 12 and 19 years (Rakesh et al., 2021). In addition, from the perspective of brain development, the age-difference of 7 years (compared to the study population in Whittle et al., 2008) is remarkable especially considering puberty, which significantly shapes brain trajectories. Brain-behaviour relationships can change with age, depending on the measurement and developmental stage, and either faster or slower neurodevelopment can be beneficial (Callaghan & Tottenham, 2016; Shaw et al., 2006; Squeglia et al., 2013). One factor that might explain variation in results is that different studies use different versions of Rothbart’s questionnaire battery, younger children typically being rated by parents, whereas older children may self-report EC. For example, Cheetham et al. (2017) and Whittle et al. (2008) both used self-reported Early Adolescent Temperament Questionnaire Revised (EATQ-R), which is a temperament and SR assessment tool for adolescents (Capaldi & Rothbart, 1992).

This study has some limitations. First, the narrow age range is both a weakness and a strength of this study. A cross-sectional study with a relatively large sample provides reliable information at this specific age-point, while a wider age range could cover different stages of development. Generally, parental report could, for example, bias the measurement towards actions that are visible to outsiders, while self-reports may be able to capture more nuanced inter-individual differences. Different questionnaire versions may also be based on slightly different items of factor structure and may differently link with neural features. Furthermore, some of the earlier studies used task-based assessment of EC (e.g. inhibitory control tasks) that may tap onto different aspects of EC compared to the parental report. Second, even though EC trait is universal and shared across cultures, the developmental outcomes (especially the ones rated using questionnaires) may vary between cultures, depending on which behaviours are valued in the society (Rothbart, 2007). This study focuses solely on Finnish 5-year-olds (many of whom from a high socioeconomic background), and the results may not generalize to other cultures and countries. Finally, the reliance on parental report is suboptimal due to its subjectivity. Parents can have biased views of their children and the parent’s own traits and abilities affect the replies. Parental report may also bias the answers towards outwardly visible behaviours, and it is subject to the parent’s own traits and abilities.

## Conclusions

Early EC and self-regulatory traits are important predictors for many social and health outcomes later in life. Therefore, it is crucial to understand neural underpinnings of EC in early childhood. In this study, we extend the prior literature by examining links between several structural brain features simultaneously (volumes, SA and CT) and child EC in typically developing 5-year-olds.

The novelty in this study comes from the finding that the regions which were significant in this study, especially the left parietal lobe, have not been linked to EC before. There are at least two possible explanations. First, the previous literature on structural cortical development of small children is so limited that the role of these regions has not been discovered before in the context of EC development. Second, this could be an age-specific finding. It is possible that the ability to self-regulate and to make plans are reflected in the parietal lobe at this stage of development, but with age other regions become increasingly important. Longitudinal studies will be crucial in exploring this possibility and to explore the developmental outcomes further.

## Acknowledgments

JJT was supported by Juho Vainio Foundation; Hospital District of Southwest Finland, Finnish State Grants for Clinical Research (ERVA / VTR); Emil Aaltonen Foundation; Alfred Kordelin Foundation; Sigrid Jusélius Foundation; Signe and Ane Gyllenberg Foundation; the Orion Research Foundation. AC was supported by Emil Aaltonen Foundation and Turku University Foundation. HM was supported by Research Council of Finland (#26080983). EK was supported by Academy of Finland (#346790), Signe and Ane Gyllenberg, Foundation, Juho Vainio Foundation and Turku University Foundation. RK was supported by The Centre for Excellence, Finnish research Council, 346121. LK was supported by Strategic Research Council (SRC) established within the Research Counsil of Finland (#352648 and subproject #352655), Signe and Ane Gyllenberg Foundation, Research Council of Finlanld #308176, Finnish State Grants for Clinical Reserach (VTR). HK was supported by Frilasarettet Eschnerska Stiftelsen. EPP was supported by Strategic Research Council (SRC) established within the Research Council of Finland (#352648 and subproject #352655), Päivikki and Sakari Sohlberg Foundation, Juho Vainio Foundation and Emil Aaltonen Foundation, Finnish Brain Foundation, Turku University Foundation, and Finnish Cultural Foundation.

We thank the FinnBrain Birth Cohort study participants, the staff, and the assisting personnel. We thank our research nurse Susanne Sinisalo, neuroradiologist Riitta Parkkola, paediatric neurologist Tuire Lähdesmäki and physicist Jani Saunavaara for their role in the 5-year MRI data collection. We thank psychologist Eeva Holmberg for her role in data collection and in the initial planning of the manuscript.

