## Supplementary Materials for "Effortful control and cortical brain structure in 5-year-old children: findings from the FinnBrain birth cohort study"

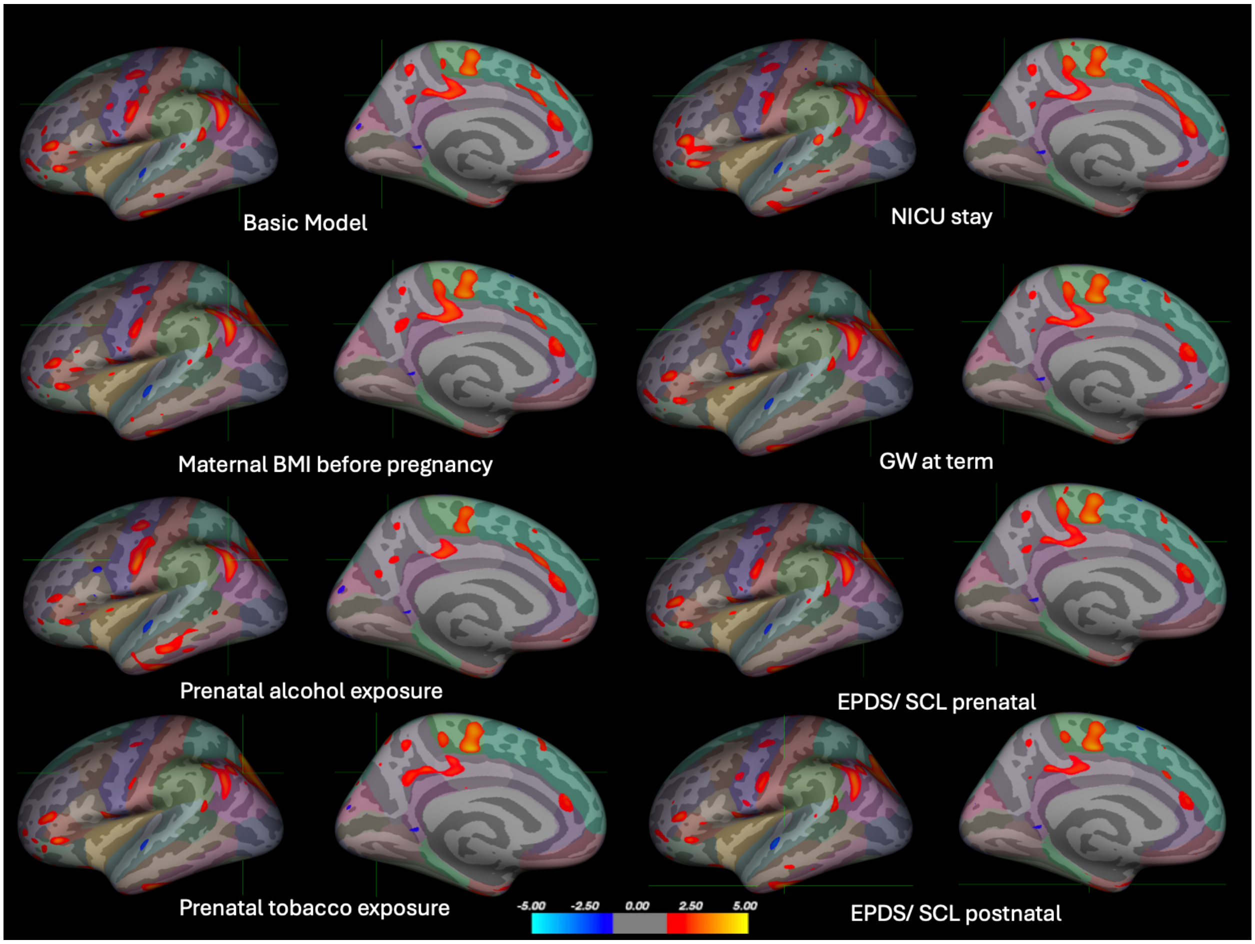


*Supplementary Figure 1: The correlation between Effortful Control (EC) and cortical volume on the left hemisphere. Covariates in all analyses were the child's gender, age at scan, ponderal index (mass in kilograms divided by height in meters cubed; measured during the neuroimaging visit), maternal age at term and maternal education level. In sensitivity analyses an additional factor was controlled for: maternal body mass index (BMI) before pregnancy, gestational weeks (GW) at term and both maternal prenatal and postnatal scores of Edinburgh Postnatal Depression Scale (EPDS) and Symptom Checklist 90 (SCL-90). In other sensitivity analyses, a part if the sample was excluded: alcohol exposure in utero with those excluded, whose mothers continued drinking after learning about pregnancy (n=134), tobacco exposure in utero, the ones with exposure excluded (n=144) and children with neonatal intensive care unit (NICU) stay excluded (n=132). Cluster color indicates significance as a z-value. Color coding of regions according to the Desikan-Killiany atlas. No correction for multiple comparisons was made.*

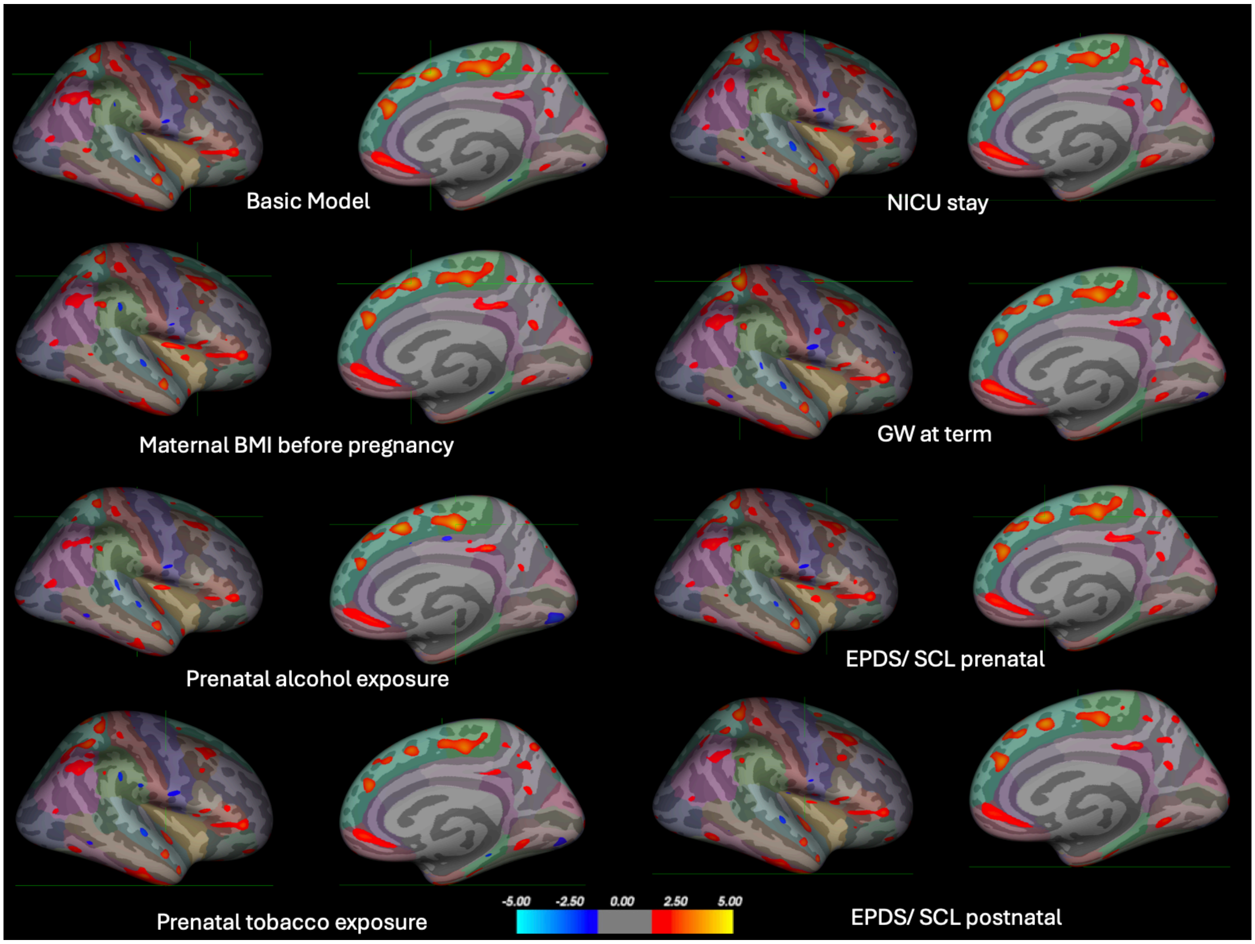


*Supplementary Figure 2: The correlation between EC and cortical volume on the right hemisphere.* *Covariates in all analyses were the child's gender, age at scan, ponderal index (mass in kilograms divided by height in meters cubed; measured during the neuroimaging visit), maternal age at term and maternal education level. In sensitivity analyses an additional factor was controlled for: maternal body mass index (BMI) before pregnancy, gestational weeks (GW) at term and both maternal prenatal and postnatal scores of Edinburgh Postnatal Depression Scale (EPDS) and Symptom Checklist 90 (SCL-90). In other sensitivity analyses, a part if the sample was excluded: alcohol exposure in utero with those excluded, whose mothers continued drinking after learning about pregnancy (n=134), tobacco exposure in utero, the ones with exposure excluded (n=144) and children with neonatal intensive care unit (NICU) stay excluded (n=132). Cluster color indicates significance as a z-value. Color coding of regions according to the Desikan-Killiany atlas. No correction for multiple comparisons was made.*

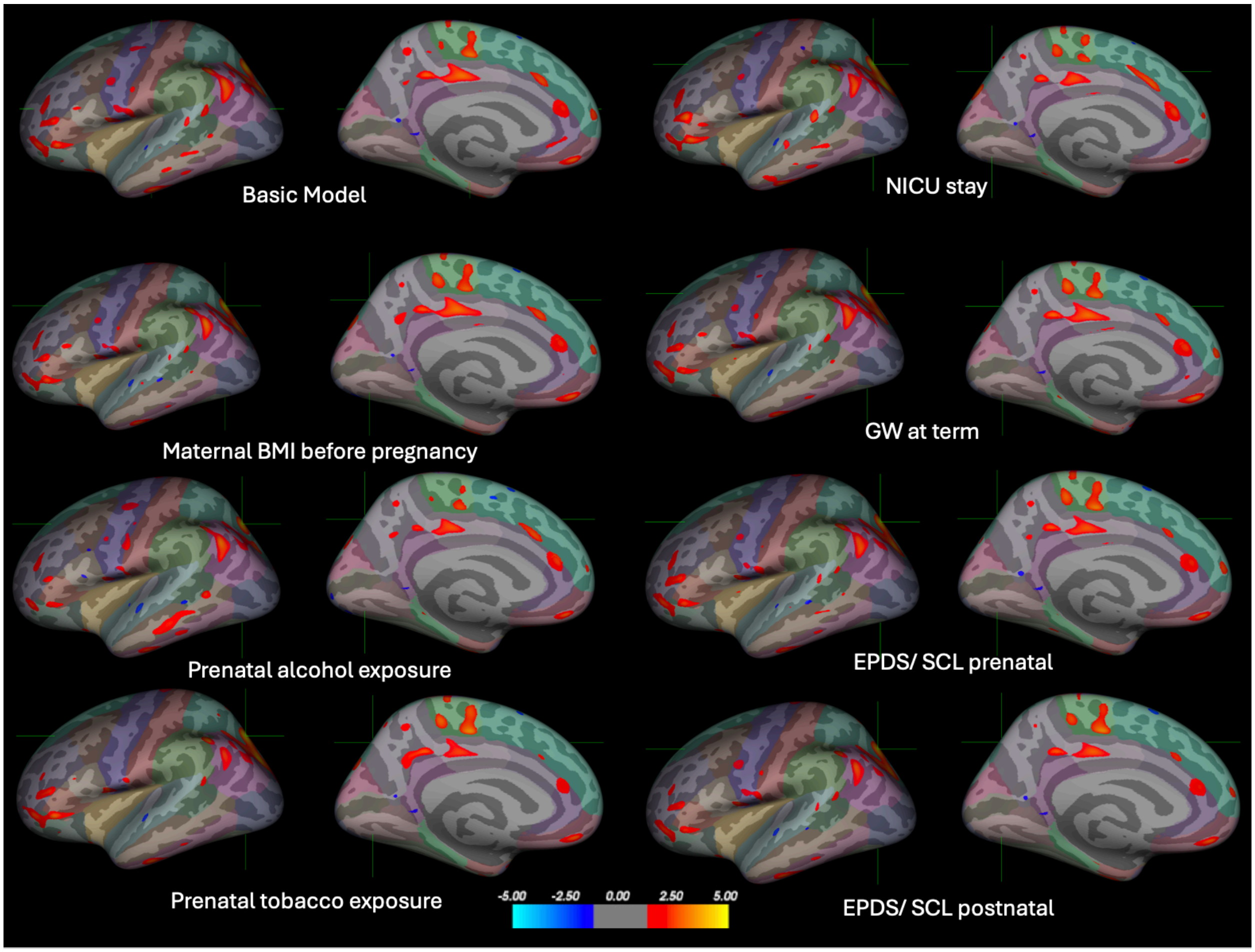


*Supplementary Figure 3: The correlation between EC and brain surface area (SA) on the left hemisphere. No significant clusters were found between EC and SA in our main results in the right hemisphere. The basic model (N=155) in the top left corner has been corrected for child's gender, age at scan, ponderal index (mass in kilograms divided by height in meters cubed; measured during the neuroimaging visit), maternal age at term and maternal education level. In sensitivity analyses an additional factor was controlled for: maternal body mass index (BMI) before pregnancy, gestational weeks (GW) at term and both maternal prenatal and postnatal scores of Edinburgh Postnatal Depression Scale (EPDS) and Symptom Checklist 90 (SCL-90). In other sensitivity analyses, a part if the sample was excluded: alcohol exposure in utero with those excluded, whose mothers continued drinking after learning about pregnancy (n=134), tobacco exposure in utero, the ones with exposure excluded (n=144) and children with neonatal intensive care unit (NICU) stay excluded (n=132). Cluster color indicates significance as a z-value. Color coding of regions according to the Desikan-Killiany atlas. No correction for multiple comparisons was made.*
