## Supplementary File 1 for "Effortful control and cortical brain structure in 5-year-old children: findings from the FinnBrain birth cohort study"

STROBE Statement—Checklist of items that should be included in reports of ***cross-sectional studies***

| | Item No | Recommendation | Page No |
|---|---|---|---|
| **Title and abstract** | 1 | (*a*) Indicate the study's design with a commonly used term in the title or the abstract | 1 |
| | | (*b*) Provide in the abstract an informative and balanced summary of what was done and what was found | 1 |
| **Introduction** | | | |
| Background/rationale | 2 | Explain the scientific background and rationale for the investigation being reported | 2 |
| Objectives | 3 | State specific objectives, including any prespecified hypotheses | Last paragraph of "Introduction" |
| **Methods** | | | |
| Study design | 4 | Present key elements of study design early in the paper | Under "Procedures" |
| Setting | 5 | Describe the setting, locations, and relevant dates, including periods of recruitment, exposure, follow-up, and data collection | Under "Participants" |
| Participants | 6 | (*a*) Give the eligibility criteria, and the sources and methods of selection of participants | Under "Participants" |
| Variables | 7 | Clearly define all outcomes, exposures, predictors, potential confounders, and effect modifiers. Give diagnostic criteria, if applicable | Under "Covariate selection for vertex wise statistics" |
| Data sources/ measurement | 8 | For each variable of interest, give sources of data and details of methods of assessment (measurement). Describe comparability of assessment methods if there is more than one group | Under "The Children's Behaviour questionnaire" and "MRI data acquisition" |
| Bias | 9 | Describe any efforts to address potential sources of bias | Under "Participants" |
| Study size | 10 | Explain how the study size was arrived at | Above Table 2 |
| Quantitative variables | 11 | Explain how quantitative variables were handled in the analyses. If applicable, describe which groupings were chosen and why | Under "Covariate selection for vertex wise statistics" |
| Statistical methods | 12 | (*a*) Describe all statistical methods, including those used to control for confounding | Under "Statistics" |
| | | (*b*) Describe any methods used to examine subgroups and interactions | NA |
| | | (*c*) Explain how missing data were addressed | Under "Neurocognitive study visit" |

| | | | |
|---|---|---|---|
| | | (*d*) If applicable, describe analytical methods taking account of sampling strategy | NA |
| | | (*e*) Describe any sensitivity analyses | Under "Covariate selection for vertex wise statistics) |
| **Results** | | | |
| Participants | 13 | (a) Report numbers of individuals at each stage of study—eg numbers potentially eligible, examined for eligibility, confirmed eligible, included in the study, completing follow-up, and analysed | Above table 2 |
| | | (b) Give reasons for non-participation at each stage | NA |
| | | (c) Consider use of a flow diagram | NA |
| Descriptive data | 14 | (a) Give characteristics of study participants (eg demographic, clinical, social) and information on exposures and potential confounders | Table 2 |
| | | (b) Indicate number of participants with missing data for each variable of interest | Table 2 |
| Outcome data | 15 | Report numbers of outcome events or summary measures | Under "Results" |
| Main results | 16 | (*a*) Give unadjusted estimates and, if applicable, confounder-adjusted estimates and their precision (eg, 95% confidence interval). Make clear which confounders were adjusted for and why they were included | Under "Covariate selection for vertex wise statistics" (some of the information not available using conventional neuroimaging tools) |
| | | (*b*) Report category boundaries when continuous variables were categorized | NA |
| | | (*c*) If relevant, consider translating estimates of relative risk into absolute risk for a meaningful time period | NA |
| Other analyses | 17 | Report other analyses done—eg analyses of subgroups and interactions, and sensitivity analyses | Under "Post Hoc analyses" |
| **Discussion** | | | |
| Key results | 18 | Summarise key results with reference to study objectives | "Discussion" paragraph 1 |
| Limitations | 19 | Discuss limitations of the study, taking into account sources of potential bias or imprecision. Discuss both direction and magnitude of any potential bias | "Discussion" last paragraph |
| Interpretation | 20 | Give a cautious overall interpretation of results considering objectives, limitations, multiplicity of analyses, results from similar studies, and other relevant evidence | Under "Discussion" |

| | | | |
|---|---|---|---|
| Generalisability | 21 | Discuss the generalisability (external validity) of the study results | Under “Conclusion” |
| **Other information** | | | |
| Funding | 22 | Give the source of funding and the role of the funders for the present study and, if applicable, for the original study on which the present article is based | Under “Acknowledgements” |

The study report follows the Strenghtening the Reporting of Observational Studies in Epidemiology (STROBE) guidelines (Vandenbroucke et al., 2007), Information on the STROBE Initiative is available at www.strobe-statement.org.
